# CLIMB (the Cloud Infrastructure for Microbial Bioinformatics): an online resource for the medical microbiology community

**DOI:** 10.1101/064451

**Authors:** Thomas R. Connor, Nicholas J. Loman, Simon Thompson, Andy Smith, Joel Southgate, Radoslaw Poplawski, Matthew J. Bull, Emily Richardson, Matthew Ismail, Simon Elwood-Thompson, Christine Kitchen, Martyn Guest, Marius Bakke, Sam K. Sheppard, Mark J. Pallen

## Abstract

The increasing availability and decreasing cost of high-throughput sequencing has transformed academic medical microbiology, delivering an explosion in available genomes while also driving advances in bioinformatics. However, many microbiologists are unable to exploit the resulting large genomics datasets because they do not have access to relevant computational resources and to an appropriate bioinformatics infrastructure. Here, we present the Cloud Infrastructure for Microbial Bioinformatics (CLIMB) facility, a shared computing infrastructure that has been designed from the ground up to provide an environment where microbiologists can share and reuse methods and data.

**DATA SUMMARY:** The paper describes a new, freely available public resource and therefore no data has been generated. The resource can be accessed at http://www.climb.ac.uk. Source code for software developed for the project can be found at http://github.com/MRC-CLIMB/

**I/We confirm all supporting data, code and protocols have been provided within the article or through supplementary data files.**

**IMPACT STATEMENT:** Technological advances mean that genome sequencing is now relatively simple, quick, and affordable. However, handling large genome datasets remains a significant challenge for many microbiologists, with substantial requirements for computational resources and expertise in data storage and analysis. This has led to fragmentary approaches to software development and data sharing that reduce the reproducibility of research and limits opportunities for bioinformatics training. Here, we describe a nationwide electronic infrastructure that has been designed to support the UK microbiology community, providing simple mechanisms for accessing large, shared, computational resources designed to meet the bioinformatic needs of microbiologists.

## INTRODUCTION

Genome sequencing has transformed the scale of questions that can be addressed by biological researchers. Since the publication of the first bacterial genome sequence over twenty years ago (1), there has been an explosion in the production of microbial genome sequence data, fuelled most recently by high-throughput sequencing (2). This has placed microbiology at the forefront of data-driven science (3). As a consequence, there is now huge demand for physical and computational infrastructures to produce, analyse and share microbiological software and datasets and a requirement for trained bioinformaticians that can use genome data to address important questions in microbiology (4). It is worth stressing that microbial genomics, with its focus on the riotous variation seen in microbial genomes, brings challenges altogether different from the analysis of the larger but less variable genomes of humans, animals or plants.

One solution to the data-deluge challenge is for every microbiology research group to establish their own dedicated bioinformatics hardware and software. However, this entails considerable upfront infrastructure costs and great inefficiencies of effort, while also encouraging a working-in-silos mentality, which makes it difficult to share data and pipelines and thus hard to replicate research. Cloud computing provides an alternative approach that facilitates the use of large genome datasets in biological research (5).

The cloud-computing approach incorporates a shared online computational infrastructure, which spares the end user from worrying about technical issues such as the installation maintenance and, even, the location of physical computing resources, together with other potentially troubling issues such as systems administration, data sharing, scalability, security and backup. At the heart of cloud computing lies *virtualization*, an approach in which a physical computing set-up is re-purposed into a scalable system of multiple independent *virtual machines*, each of which can be pre-loaded with software, customised by end users and saved as *snapshots* for re-use by others. Ideally, such an infrastructure also provides large-scale data storage and compute capacity on demand, reducing costs to the public purse by optimising utilization of hardware and avoiding resources sitting idle while still capitalising on the economies of scale.

The potential for cloud computing in biological research has been recognized by funding organisations and has seen the development of nationwide resources such as iPlant (6) (now CyVerse), NECTAR (7) and XSEDE (8) that provide researchers with access to large cloud infrastructures. Here, we describe a new facility, designed specifically for microbiologists, to provide a computational and bioinformatics infrastructure for the UK’s academic medical microbiology community, facilitating training, access to hardware and sharing of data and software.

## Resource overview

The Cloud Infrastructure for Microbial Bioinformatics (CLIMB) facility is intended as a general solution to pressing issues in big-data microbiology. The resource comprises a core physical infrastructure (Figure 1), combined with three key features making the cloud suitable for microbiologists.

**Figure 1.**
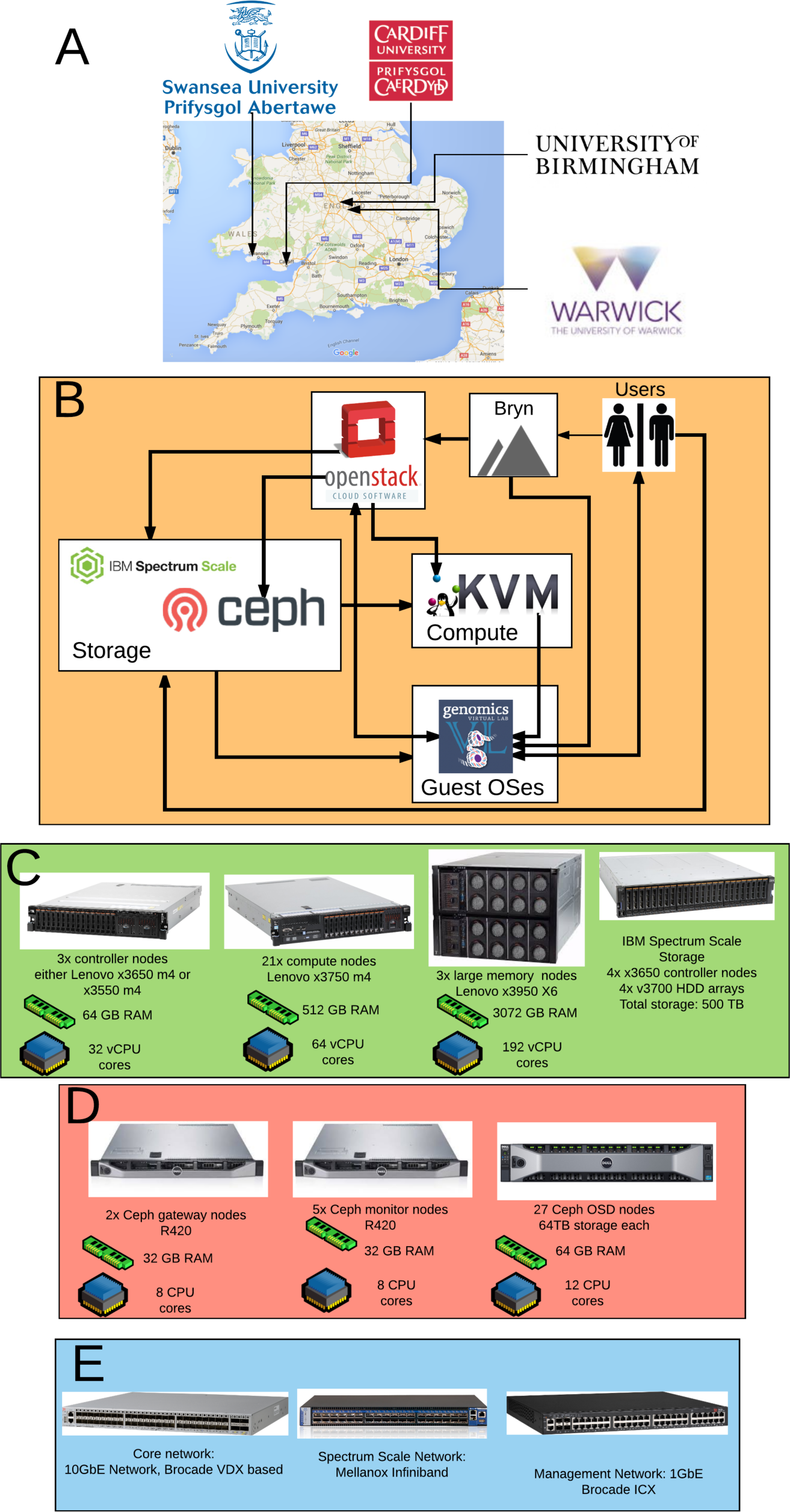
Overview of the system. A. The sites where the computational hardware is based. B. High level overview of the system and how the different software components connect to one another. C. Compute hardware present at each of the four sites. D. Hardware comprising the Ceph storage system at each site. E. Type and role of network hardware used at each site.

First, CLIMB provides a single environment, complete with pipelines and datasets that are co-located with computing resource. This makes the process of accessing published packages and sequence data simpler and faster, improving reuse of software and data.

Second, CLIMB has been designed with training in mind. Rather than having trainees configure personal laptops or face challenges in gaining access to shared high performance computing resources, we provide training images on virtual machines that have all the necessary software installed and we provide each trainee with her own personal server to continue learning after the workshop concludes.

Third, by bringing together expert bioinformaticians, educators and biologists in a unified system, CLIMB provides an environment where researchers across institutions can share data and code, permitting complex projects iteratively to be remixed, reproduced, updated and improved.

The CLIMB core infrastructure is a cloud system running the free open-source cloud operating system OpenStack (9). This system allows us to run over 1000 virtual machines at any one time, each preconfigured with a standard user configuration. Across the cloud, we have access to almost 43 terabytes of RAM. Specialist users can request access to one of our twelve high-memory virtual machines each with 3 terabytes of RAM for especially large, complex analyses (Figure 1). The system is spread over four sites to enhance its resilience and is supported by local scratch storage of 500TB per site employing IBM’s Spectrum Scale storage (formerly GPFS). The system is underpinned by a large shared object storage system that provides approximately 2.5 petabytes of data storage, which may be replicated between sites. This storage system, running the free open-source Ceph system (10), provides a place to store and share very large microbial datasets—for comparison, the bacterial component of the European Nucleotide Archive is currently around 400 terabytes in size. The CLIMB system can be coupled to sequencing services; for example, sequence data generated by the MicrobesNG service (11) has been made available within the CLIMB system.

## Resource performance

To assess the performance of the CLIMB system in comparison to traditional High Performance Computing (HPC) systems and similar cloud systems, we undertook a small-scale benchmarking exercise (Figure 2). Compared to the Raven HPC resource at Cardiff (running Intel processors a generation behind those in CLIMB), performance on CLIMB was generally good, offering a relative increase in performance of up to 38% on tasks commonly undertaken by microbial bioinformaticians. The CLIMB system also compares well to cloud servers from major providers, offering better aggregate performance than Microsoft Azure A8 and Google N1S2 virtual machines. The results also reveal a number of features that may be relevant to where a user chooses to run their analysis. CLIMB performs worse than Raven when running BEAST, and provides a limited increase in performance for the package nhmmer, suggesting that while it is possible to run these analyses on CLIMB, other resources–such as local HPC facilities might be more appropriate. Conversely, the largest performance increases are observed for Prokka, Snippy and PhyML, which encompass some of the most commonly used analyses undertaken in microbial genomics. It is also interesting to note that both commercial clouds offer excellent performance relative to Raven for two workloads; muscle and PhyML. The source of this performance is difficult to predict, but it is possible that these workloads may be more similar to the sort of workloads that these cloud services have been designed to handle. On the basis of the performance results more generally, however, CLIMB is likely to offer a number of performance benefits over local resources for many microbial bioinformatics workloads.

**Figure 2.**
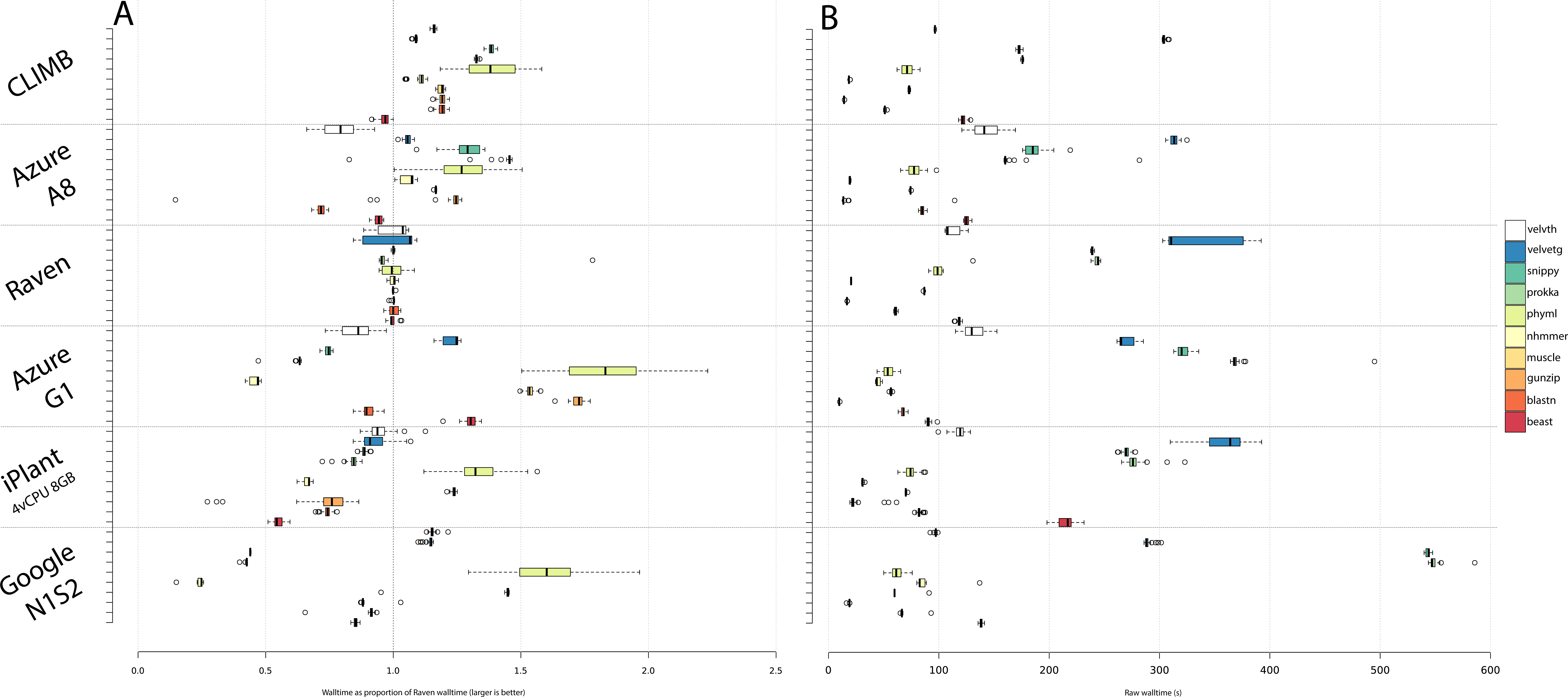
(appended at end of document). Relative performance of VMs running on cloud services, compared to the Cardiff University HPC system, Raven. A. Values for each package are the mean of the wall time taken for 10 runs performed on Raven, divided by the mean wall time of 40 runs undertaken on the VM on the named service. Values greater than 1 are faster than Raven, values less than 1 are slower. B. Showing the raw wall time values for the named software on each of the systems.

## Providing a single environment for training, data and software sharing

The CLIMB system is accessed through the Internet, via a simple set of web interfaces enabling the sharing of software on virtual machines (Figure 3). Users request a virtual machine via a web form. Each virtual machine makes available the microbial version of the Genomics Virtual Laboratory (Figure 3) (7). This includes a set of web tools (Galaxy, Jupyter Notebook and RStudio, with an optional PacBio SMRT portal), as well as a set of pipelines and tools that can be accessed via the command line. This standardised environment provides a common platform for teaching, while the base image provides a versatile platform that can be customized to meet the needs of individual researchers.

**Figure 3.**
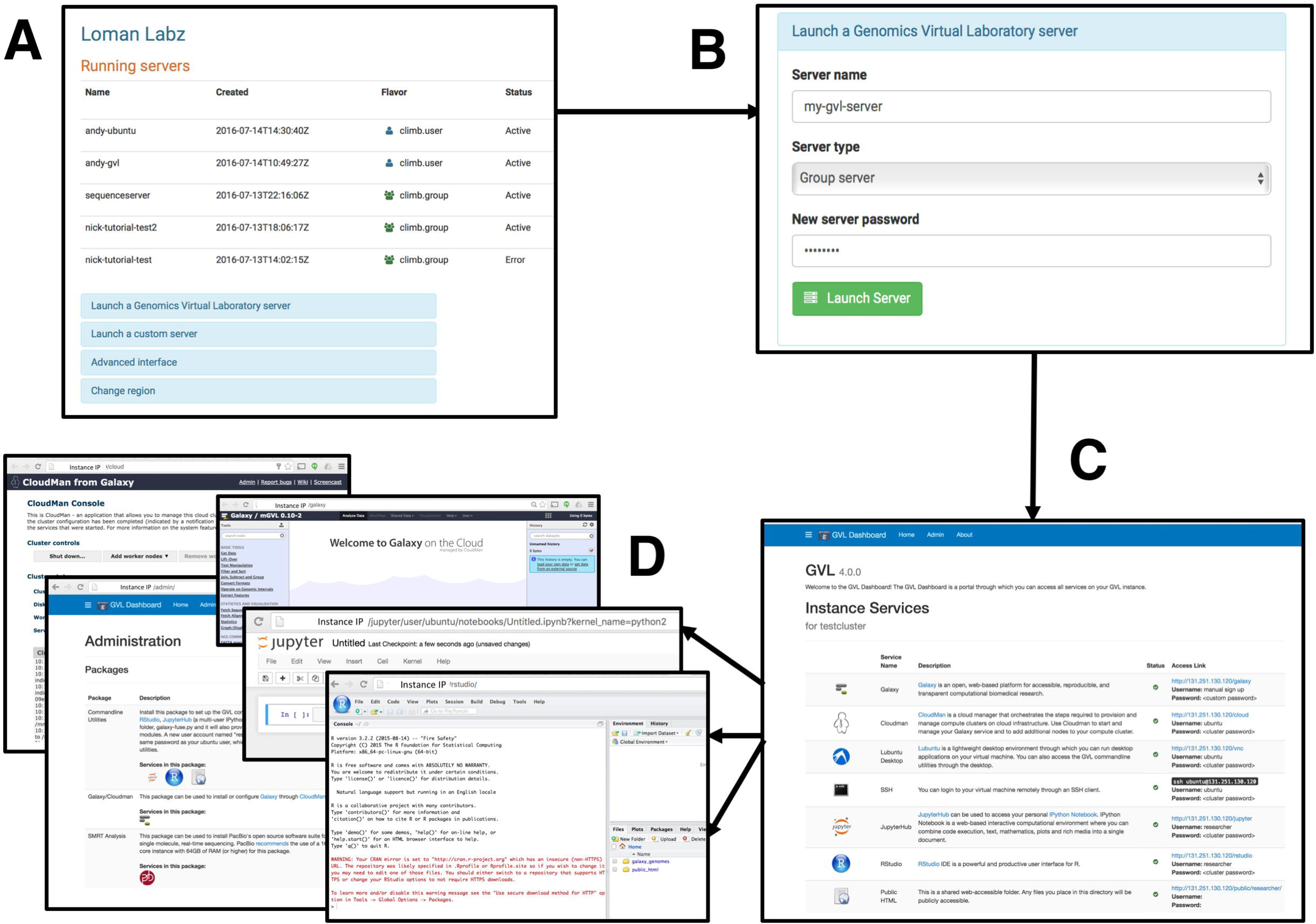
(appended at end of document) CLIMB virtual machine launch workflow. A user, on logging in to the Bryn launcher interface is presented with a list of the virtual machines they are running and are able to stop, reboot or terminate them (panel A). Users launch a new GVL server with a minimal interface, specifying a name, the server ‘flavour’ (user or group) and an access password (panel B). On booting, the user accesses a webserver running on the GVL instance, which gives access to various services that are started automatically (panel C). The GVL provides access to a Cloudman, a Galaxy server, an administration interface, Jupyter notebook and RStudio (panel D, top to bottom).

## System access

Users register at our website, using their UK academic credentials (http://bryn.climb.ac.uk/). Upon registration, users have one of two modes of access: the first is to launch an instance running a preconfigured virtual machine, with a set of predefined pipelines or tools, which includes the Genomics Virtual Laboratory. The second option is aimed at expert bioinformaticians and developers who may want to be able to develop their own virtual machines from a base image—to enable this we also allow users to access the system via a dashboard, similar to that provided by Amazon Web Services, where users can specify the size and type of virtual machine that they would like, with the system then provisioning this up on demand. To share the resource fairly, users will have individual quotas that can be increased on request. Irrespective of quota size, access to the system is free of charge to UK academic users.

## CONCLUSION

CLIMB is probably the largest computer system dedicated to microbiology in the world. The system has already been used to address microbiological questions featuring bacteria (12) and viruses (13). CLIMB has been designed from the ground up to meet the needs of microbiologists, providing a core infrastructure that is presented in a simple, intuitive way. Individual elements of the system—such as the large shared storage and extremely large memory systems—provide capabilities that are usually not available locally to microbiologists within most UK institutions, while the shared nature of the system provides new opportunities for data and software sharing that have the potential enhance research reproducibility in data intensive biology. Cloud computing clearly has the potential to revolutionize how biological data are shared and analysed. We hope that the microbiology research community will capitalise on these new opportunities by exploiting the CLIMB facility.

## DATA BIBLIOGRAPHY

Not applicable

## Supplementary Table 1.

Table contains the raw wall time recorded for 40 independent runs on each of the indicated comparison systems. For figure 2 these raw data were compared against the mean wall time recorded from 10 runs on the Cardiff University Raven system. The raw and calculated values are in the attached spreadsheet, in different workbooks.

## ACKNOWLEDGEMENTS

We would like to especially acknowledge the assistance of Simon Gladman, Andrew Lonie, Torsten Seeman and Nuwan Goonasekera for their extensive assistance in getting the GVL running on CLIMB. We thank Isabel Dodd (Warwick) and Ben Pascoe (Bath) for assistance managing the project, and would also like to thank the local University network and IT staff (particularly Dr Ian Merrick and Kevin Munn at Cardiff and Chris Jones at Swansea) who have helped to get the system up and running, and University procurement staff (especially Anthony Hale at Cardiff) who worked to get the system purchased in challenging timescales. In addition, we would also like to thank early access CLIMB users for testing and reporting issues with our service, with particular thanks to Phil Ashton (Oxford University Clinical Research Unit Vietnam), Ed Feil, Harry Thorpe, Sion Bayliss (University of Bath). We thank Emily Richardson (University of Birmingham) for developing tutorial materials for CLIMB and testing. We thank all our project partners and suppliers at OCF, IBM, Red Hat/Ceph, Dell, Mellanox and Brocade for support of the project with particular thanks to Arif Ali and Georgina Ellis (OCF), Dave Coughlin and Henry Bennett (Dell), Ben Harrison (RedHat), Stephan Hohn (RedHat/InkTank), Jim Roche (Lenovo).

